# PlantConnectome: knowledge graph encompassing >70,000 plant articles

**DOI:** 10.1101/2023.07.11.548541

**Authors:** Shan Chun Lim, Kevin Fo, Rohan Shawn Sunil, Manoj Itharajula, Yu Song Chuah, Herman Foo, Emilia Emmanuelle Davey, Melissa Fullwood, Guillaume Thibault, Marek Mutwil

## Abstract

One of the main quests of plant biology is understanding how genes and metabolites work together to form complex networks that drive plant growth, development, and responses to environmental stimuli. However, the ever-growing volume and diversity of scientific literature make it increasingly challenging to stay current with the latest advances in gene function studies. Here, we tackle the challenge by deploying the text-mining capacities of large language models to process over 71,000 plant biology abstracts. Our approach unveiled nearly 5 million functional relationships between a wide array of biological entities—genes, metabolites, tissues, and others—with a high accuracy of over 85%. We encapsulated these findings in PlantConnectome, a user-friendly database, and demonstrated its diverse utility by providing insights into gene regulatory networks, protein-protein interactions, and stress responses. We believe this innovative use of AI in the life sciences will allow plant scientists to keep up to date with the rapidly growing corpus of scientific literature. PlantConnectome is available at https://plant.connectome.tools/.

## Introduction

Despite decades of research, only ∼15% of *Arabidopsis thaliana*’s genes have been comprehensively characterized, and the rate of new articles reporting gene functions has dropped to <30% in 2023 since the peak in 2008 (Sunil et al. 2024). Due to the time-consuming experiments and increased requirements to publish in premier journals, the time needed to characterize a gene can take several years. Thus, choosing which gene to start characterizing requires a strong hypothesis, which is typically based on previous work reported in the literature. However, staying up to date with the continuously growing scientific literature, and integrating the numerous pieces of the gene function puzzle can be time-consuming and limit our ability to form strong hypotheses.

Alternatively, computational gene function prediction can suggest which genes have a specific function and are invaluable in generating new gene function hypotheses (Brown et al. 2005; Persson et al. 2005). Predicting gene function requires two components: i) omics data that captures gene properties (e.g., coding sequence, expression patterns, and protein structure) and ii), gold standard data (i.e., genes with experimentally verified functions) (Radivojac et al. 2013; Rhee and Mutwil 2014). The omics data is firstly used to connect uncharacterized and characterized genes based on sequence or expression similarity. Then, uncharacterized genes are labeled according to the functions of the characterized genes (i.e., the gold standard data) to which they were connected (Rhee and Mutwil 2014).

Nonetheless, gene function prediction remains highly challenging due to the complexity and vastness of biological data, plateauing our understanding of gene functions (Radivojac et al. 2013). Specifically, establishing the gold standard necessitates manual, work-intensive extraction of gene functional information from scientific articles (Oughtred et al. 2021), preventing public repositories that harbor the gold standard data, such as BioGRID (protein-protein interactions, or PPIs) and AGRIS (gene regulatory networks, or GRNs)(Yilmaz et al. 2011; Oughtred et al. 2021), from keeping up to date with state-of-the-art knowledge. Furthermore, such repositories are typically restricted to specific data types (e.g., PPI or GRNs), precluding the integration of various data kinds that is critical to deepening our understanding of plant biology.

Several methods that extract gene functional information from literature have been developed to address these challenges. PL-PPF (Predicate Logic for Predicting Protein Functions) uses statistical methods to infer if a protein and a molecular term that describes protein function are semantically related (Taha et al. 2019). However, the method requires constructing complex statistical and linguistic models to link protein to function and only considers protein-function relationships. The EVEX database processes abstracts and full texts to identify regulatory relationships, posttranslational modifications, gene expression patterns and other features of genes (Landeghem et al. 2013). However, the method also requires a manually constructed complex set of rules to extract and categorise the relationships, and the database has not been updated for a while. Another approach uses non-negative matrix factorization (NMF) for feature reduction and then classifies the function of genes using K-nearest neighbor (KNN)(Fodeh and Tiwari 2018). While the approach can reveal gene functions (e.g., gene A is a transcription factor), it does not reveal gene-gene relationships (e.g., gene A regulates gene B). STRING is a popular database that integrates protein-protein data, genomic features, co-expression and text mining to build gene co-function networks (Szklarczyk et al. 2023). However, the text mining approach only identifies genes that are frequently mentioned together and cannot reveal the type of the relationship (e.g., interaction, regulation, activation) or identify relationships between genes and other entities (e.g., treatments, hormones). KnetMiner uses a rule-based approach to build knowledge graphs capturing relationships between various entities (Hassani-Pak et al. 2016). However, the rule-based system requires the integration of multiple heterogenous datasets (e.g., 12 types of data)(Hassani-Pak et al. 2016), making it difficult to include new data, species and evidence types into the network. Furthermore, the database is gene-centric and does not allow searching the knowledge graph to understand how the different types of entities are related (e.g., traits and hormones).

In this paper, we aim to address the two fundamental challenges in gene function prediction: integrating the burgeoning information from scientific literature and using it to generate gold standard data for gene function prediction approaches. To achieve this, we seized the recent developments in Large Language Models (LLMs) to process over 71,000 research papers from leading journals in plant biology. Our approach excavated 4.8 million functional relationships between more than 2.7 million entities comprising genes, metabolites, tissues, organs, and other biological components. The manual inspection of these relationships revealed not only their high accuracy but also their ability to identify functional relationships between biological entities, even doubling the amount of functional information relative to the current coverage of gene regulatory networks. To provide access to this data, we constructed PlantConnectome, a user-friendly database containing knowledge graphs that can illuminate gene function, organ development, gene regulatory networks, protein-protein interactions, and other biological entities. PlantConnectome is available at https://plant.connectome.tools/.

## Materials and Methods

### Retrieval of articles

Using BioPython version 1.81, we downloaded papers containing *Arabidopsis thaliana* species name and gene identifiers (e.g., *At4g32410*). For each gene identifier, we also searched with gene aliases (e.g., *CESA1, RSW1*) retrieved from www.arabidopsis.org (Table S1). The NCBI query was: *query = f’(Arabidopsis thaliana[Title/Abstract] AND {query_term}[tw]) OR (Arabidopsis[Title/Abstract] AND {query_term}[tw]) OR (Thale cress[Title/Abstract] AND {query_term}[tw] OR (Mouse ear cress[Title/Abstract] AND {query_term}[tw] OR (Mouse-ear cress[Title/Abstract] AND {query_term}[tw])’.* Query_term are the genes in Table S1, and other alternative names of Arabidopsis were included in the search. The code to perform this analysis is available in the Colab Notebook in Supplementary Data 1.

### Large language model analysis of articles

We used GPT4 models to extract entities and relationships (GPT-4o), entity definitions (GPT-4o-mini) and to identify the species that the article uses as a model (GPT-4o). Furthermore, for each entity relationship, we asked GPT-4o to identify the evidence (e.g., yeast two-hybrid, bioinformatical prediction) underpinning the relationship. We iterated over several prompts to arrive at prompts that yielded consistently accurate results on selected papers, such as Brinngmann et al., 2012, and others. In total, **71,136** articles were processed using OpenAI’s batch API (Application Programming Interface)(https://openai.com/api/pricing/). The code to perform this analysis is available as Colab Notebook in Supplementary Data 1.

### Entity and relationship disambiguation

To disambiguate relationships (e.g., ‘caused’, ‘cause’, ‘causes’) and entities (e.g., ‘Arabidopsis plants’, ‘Arabidopsis’, ‘Arabidopsis thaliana’), we identified the top 100 most common relationships and entities and devised a rule-based method to map the various synonyms or variations to a canonical form. Passive edges (e.g., ‘is regulated by’) were converted to active form (‘regulates’). Entities that differed by casing (e.g., ‘Genes’, ‘genes’) were represented by one canonical form. The code used for this section is available as Supplementary Data 1.

### Construction of PlantConnectome database

The PlantConnectome is hosted on a Google Cloud server. The backend was implemented using the Python framework Flask and the Python packages networkx version 3.1, pickle version 3.11.4, json version 3.11.4, and regex *version 3.11.4*. We used JavaScript dependencies jQuery v3.6, Cytoscape.js v3.23, ChartJS v4.3, and FileSaver v2.0.5 to visualize the knowledge graphs. The GitHub repository containing the source code of the database is available at https://github.com/mutwil/plant_connectome_latest.git

### API for PlantConnectome

PlantConnectome also has an application programming interface that allows users to conduct search queries remotely. The API accepts GET requests and is implemented using the same set of packages described earlier. For each successful call to PlantConnectome’s API, a JSON (JavaScript Object Notation) object is returned, containing the functional abbreviations, GO terms, other nodes, and text summaries associated with the search query. To perform searches using the API, users can add “/api/<search type>/<search query>” to the web address, where “<search type=” and “<search query>” are placeholders representing the type of search and user’s query, respectively.

## Results

### Meta-analysis of 71,136 Paper Abstracts

To retrieve articles that focus on *Arabidopsis thaliana* genes and how these genes are related to other biological entities, we searched for articles that mention *Arabidopsis thaliana* and gene IDs in the abstracts (code available in Supplementary Data 1, gene IDs Table S1). In total, 71,136 articles, of which 19,809 and 51,327 were accessible as full-text articles or abstracts only, respectively (Figure 1A, Table S2). The top 20 journals comprise Plant Physiology, the Plant Journal and Plant Cell, for which most articles were not available for high-throughput download as full text (Figure 1A, red bars). Conversely, the open access policies and the option to programmatically download the articles of the Frontiers in Plant Science, PLOS One, New Phytologist, BMC Plant Biology and Scientific Reports allowed us to download full-text articles from these journals.

**Figure 1.**
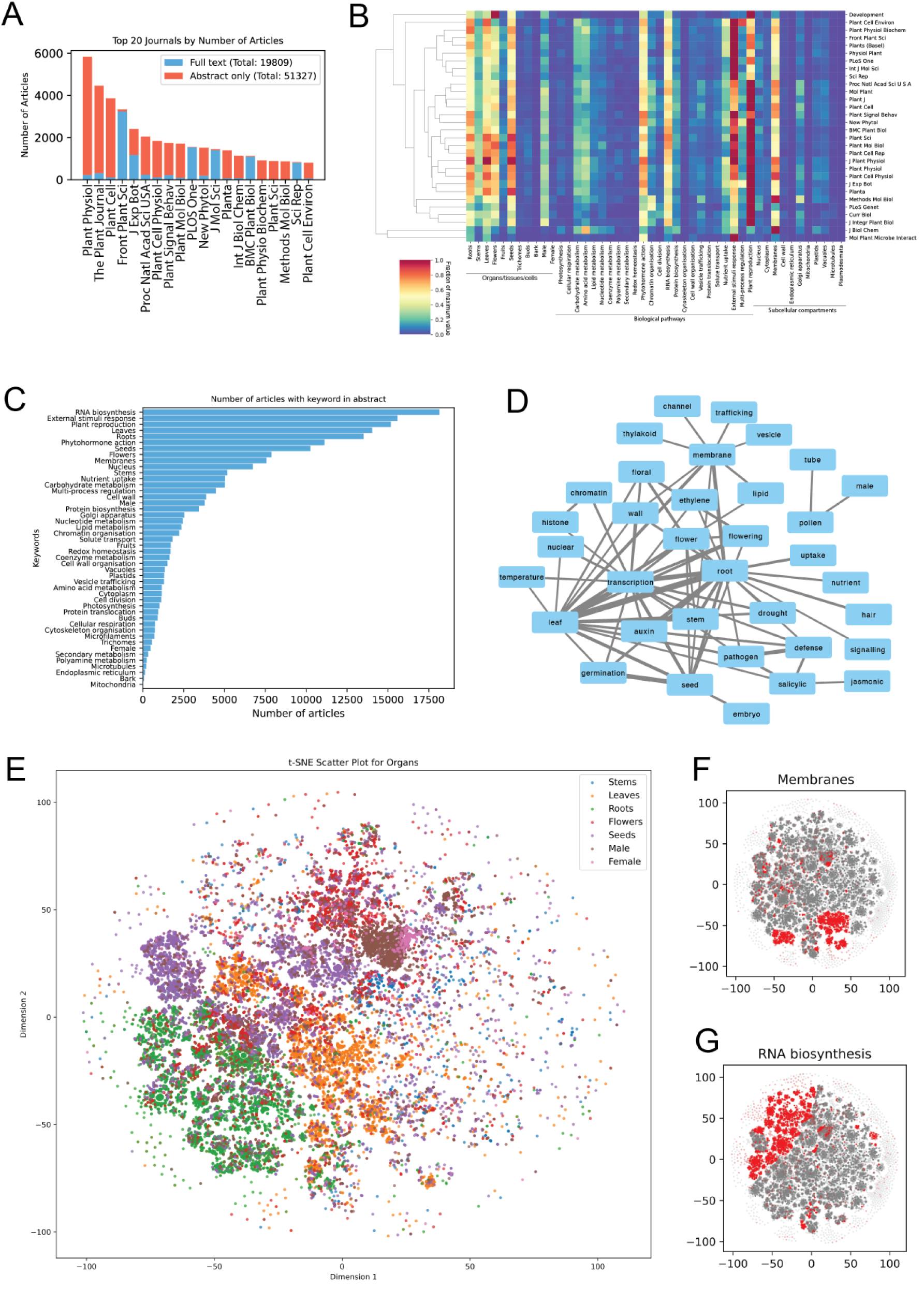
Meta-analysis of the 71,136 article abstracts. Meta-analysis of plant literature. A) Top 20 journals of the 71,136 articles analyzed in this study. The red and blue bars indicate the abstract-only or full-text articles, respectively. B) Clustermap of top 30 journals (rows) and topics (columns). The colormap corresponds to the fraction of a maximum value found in a row (journal) across the 71,136 articles. C) The number of abstracts (x-axis) with a given keyword (y-axis) of the analyzed papers. Each abstract can contain multiple keywords. D) Co-occurrence network of keywords in abstracts. Nodes represent keywords, while edges connect keywords found in at least 600 abstracts. Edge width is proportional to the number of abstract where any two keywords are co-mentioned. E) t-SNE visualization of the abstracts with a focus on plant organs. Each point represents an article, and the colors indicate the different organs. The plots for F) ‘membranes’ and G) ‘RNA biosynthesis’ are shown to the right.

To investigate whether the top 20 journals tend to publish specific topics, we determined the surveyed journals’ discussion of cellular compartments, organs, and biological functions to assess their considered research topics. To this end, we defined a list of keywords pertaining to organs (e.g., roots = [root, hair, nodule, mycorrhizae]), biological processes (photosynthesis = [photosynthesis, photorespiration, photosystem]), and cellular compartments (e.g., nucleus = [nucleus, nucleolus, chromosome, nuclear pore])(Table S3). We counted the number of these keywords in each abstract (Supplementary Data 2). Most journals did not show particular specificity for any topic, except Development (focus on reproduction, red cell), Journal of Biological Chemistry (membranes) and Molecular Plant-Microbe Interactions (external stimuli responses)(Figure 1B-C). The most commonly studied organs were: roots, leaves, flowers and seeds, and the most studied pathways were: phytohormone action (how hormones work), external stimuli response (how plants respond to the environment), RNA biosynthesis (how gene expression is regulated) and plant reproduction and most studied subcellular compartments were: membranes, and Golgi apparatus (Figure 1B-C). Next, we investigated which keywords tend to co-occur in abstracts (Table S4, Supplementary Data 3), which revealed, e.g., that pathogen research focuses on salicylic acid and transcriptional responses and uses leaves and roots as model organs (Figure 1D).

To visualize the relationships among the abstracts, we generated a two-dimensional t-distributed Stochastic Neighbor Embedding (t-SNE) plot (Macosko et al. 2015), of the keyword counts (Supplementary Data 2). This technique allows us to represent high-dimensional data in a way that preserves local similarities between abstracts. We used a perplexity value of 40, which balances the attention between local and global aspects of the data, and ran 1,000 iterations to ensure convergence to a stable configuration (Figure S1 shows the influence of t-SNE parameters). The resulting plot provided an interpretable layout that highlighted clusters of abstracts with similar content or themes. The plots demonstrate clear groupings by biological processes (Figure S2), subcellular compartments (Figure S3) and organs (Figure S4), providing a bird’s eye view of plant literature (Figure 1E).

### Text Mining Research Papers with Large Language Model Reveals 4,819,469 Relationships between 2,771,008 Entities

To extract information pertaining genes, metabolites, organs, environmental conditions, and other entities, we tasked OpenAI’s GPT models with identifying functional relationships between pairs of entities (e.g., ‘gene A’ -interacts with - ‘gene B’)(Figure 2A, prompt 1 with GPT-4o), and also identifying the types of each entity (genes, metabolites, organs, treatments, others)(Supplementary Data 1). The output of this analysis was a Knowledge Graph (KG), where nodes represent entities and edges represent relationships (e.g., ‘interacts with’, ‘regulates’, ‘causes’). To better understand which types of evidence underpin each relationship (e.g., ‘pull-down assay’, ‘co-expression analysis’), we also asked GPT-4o model to reveal the relationship basis and species the experiments were performed in (Figure 2A, prompt 2 with GPT-4o). Finally, we tasked GPT-4o-mini model to annotate the extracted entities (e.g., ‘CESA’ - is - ‘Cellulose Synthase A’)(Figure 2A, prompt 3 with GPT-4o-mini). The process yielded a large KG comprising 4,819,469 relationships between 2,771,008 entities (Supplementary Data 4, Table S5 contains the three prompts).

**Figure 2.**
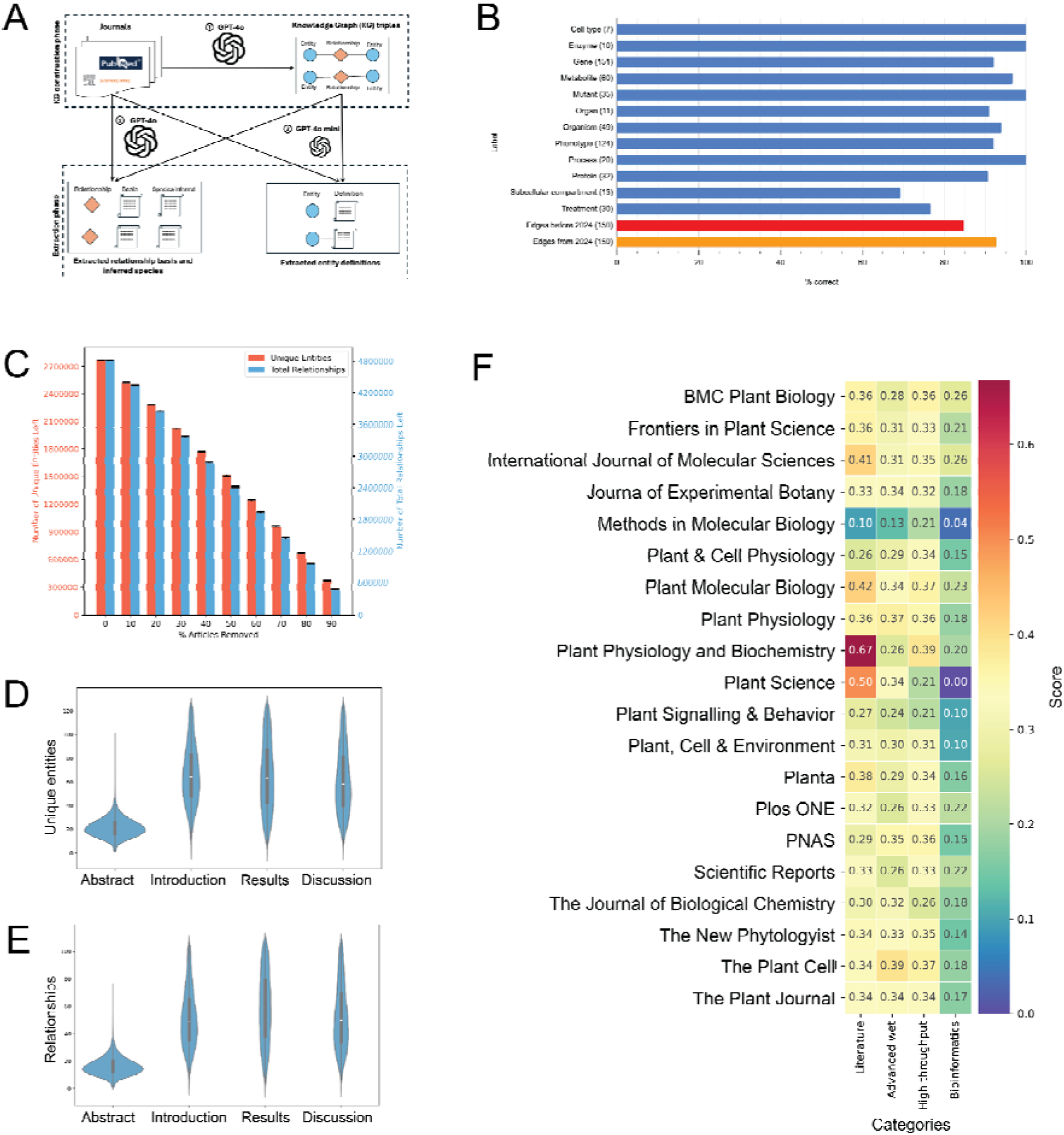
Evaluation of the plant knowledge graph. A) The pipeline to extract: 1. the knowledge graph from the literature, 2. species and relationship basis and 3. entity definitions. B) The percentage of correct entities (blue bars) and relationships (red and orange bars). The x-axis indicates the percentage of correct items inferred from manual curation. C) The number of unique entities (red bars, left y-axis) and relationships (i.e., number of edges, blue bars, right y-axis) as a function of % articles removed (x-axis). The error bars represent the standard deviation. The data was generated by randomly removing a given percentage of articles 100 times. D) and E) the number of relationships and entities extracted from abstracts, introductions, results and discussions, respectively. F) Average score depicting the diversity of methods extracted from the top 20 journals (Figure 1A). The method categories include ‘literature’ citations, ‘advanced wet’ lab techniques, ‘high throughput’ experiments and ‘bioinformatics’. A score of 1 indicates that a given journal has, on average, used all types of methods under one category, while a score of 0 indicates that no methods have been used. A method category is comprised of keywords. For example, ‘bioinformatics’ is comprised of: ‘sequence’, ‘phylogenetic’, ‘genomic’, ‘alignment’, and ‘differential expression’. The code and keywords are available in Supplementary Data 4.

Large language models are known to hallucinate and misunderstand the text, and to evaluate the accuracy of the identified entity types, we randomly selected 300 edges from the KG (code for random selection in Supplementary Data 1). We manually evaluated whether the identified entity types (e.g., ‘flavonol’ type is ‘metabolite’) are correct, by comparing the entity types with their known biological function. Overall, we observed >90% accuracy in entity type classification, with the exception of ‘subcellular compartment’ and ‘treatment’ (Figure 2B, Table S6). The incorrectly classified ‘subcellular compartment’ entity types comprised genomic features such as ‘distal enhancer elements’, and protein domains (‘13 TM helices’)(Table S6), while incorrect ‘treatment’ types comprised methods and resources (e.g., ‘microsomal preparations’, ‘Genbank’).

To evaluate the accuracy of the edges, we manually compared them to the text from which they were extracted. Furthermore, since GPT-4 models have an October 2023 knowledge cutoff and could have been trained on the analysed articles, we chose edges from articles published before 2024 and from 2024. Overall, we observed a high accuracy of 85% (before 2024, Figure 2B red bar) and 93% (2024, orange bar), showing that the models can extract information from any text and not just regurgitate training data. The incorrect relationships typically misunderstood a hypothesis of the authors (source sentence: “(results) made us wonder whether this tissue-specific polarization of PINs is conserved in other bryophytes”), incorrectly producing a fact edge (‘tissue-specific polarization of PINs conserved in other bryophytes’)(Table S6). Since the analyzed articles were shortlisted by term ‘*Arabidopsis thaliana*’, the majority of the relationships were identified in the model plant, but we also identified other models and crops, such as rice, wheat, soybean, tobacco and even yeast (Figure S5).

To investigate the relationship between the number of articles and the number of identified entities and relationships, we randomly removed 10-90% of articles 100 times, and recounted the number of retrieved items. Overall, we observed a linear relationship between the number of articles and the retrieved data (Figure 2C), indicating that more articles would expand the KG further. The amount of information extracted from the introduction (median 50 and 63 relationships and entities extracted, Figure 2D-E), results (59, 63) and discussion (47, 57) was higher than from abstracts (16, 21). Thus, increasing the number of full text articles would further expand the KG.

Finally, we investigated which types of evidence are present in the top 20 journals. We categorized the evidence into ‘literature’ (article citing findings from other articles), ‘advanced wet’ (article using advanced experimental approaches, such as pull-down, transgenic lines), ‘high throughput’ (evidence based on, e.g., RNA-seq analysis, differential gene expression) and ‘bioinformatics’ (evidence based on sequence alignment, phylogenetic tree, Supplementary Data 1 contains the used code and keywords). Overall, the evidence profiles of the different journals were similar, with Plant Cell and Plant Physiology on average using more advanced wet lab methods (advanced wet > 0.35). Plant Science, Plant Physiology and Biochemistry contain extensive literature-based evidence. In contrast, Methods in Molecular Biology which focuses typically on one method, contained the least diversity of the used evidence types.

### Properties of the Connectome Network

We used the KG to construct the Connectome network, and visual inspection of the whole graph revealed cluster-like structures of densely connected entities (Figure 3A). Certain networks, such as protein-protein interactions, display scale-free behavior, where most nodes have few connections, and few nodes have many connections (Broido and Clauset 2019). To investigate whether the Connectome is scale-free, we constructed a scatterplot of its log-transformed node frequency (*p(k)*) and node degree (*k*)(Figure 3B). The points formed a line with a negative slope, indicating a typical power law distribution (Mutwil et al. 2010), indicating that most entities have a few relationships, while a small number of entities act as hubs with a large number of connections.

**Figure 3.**
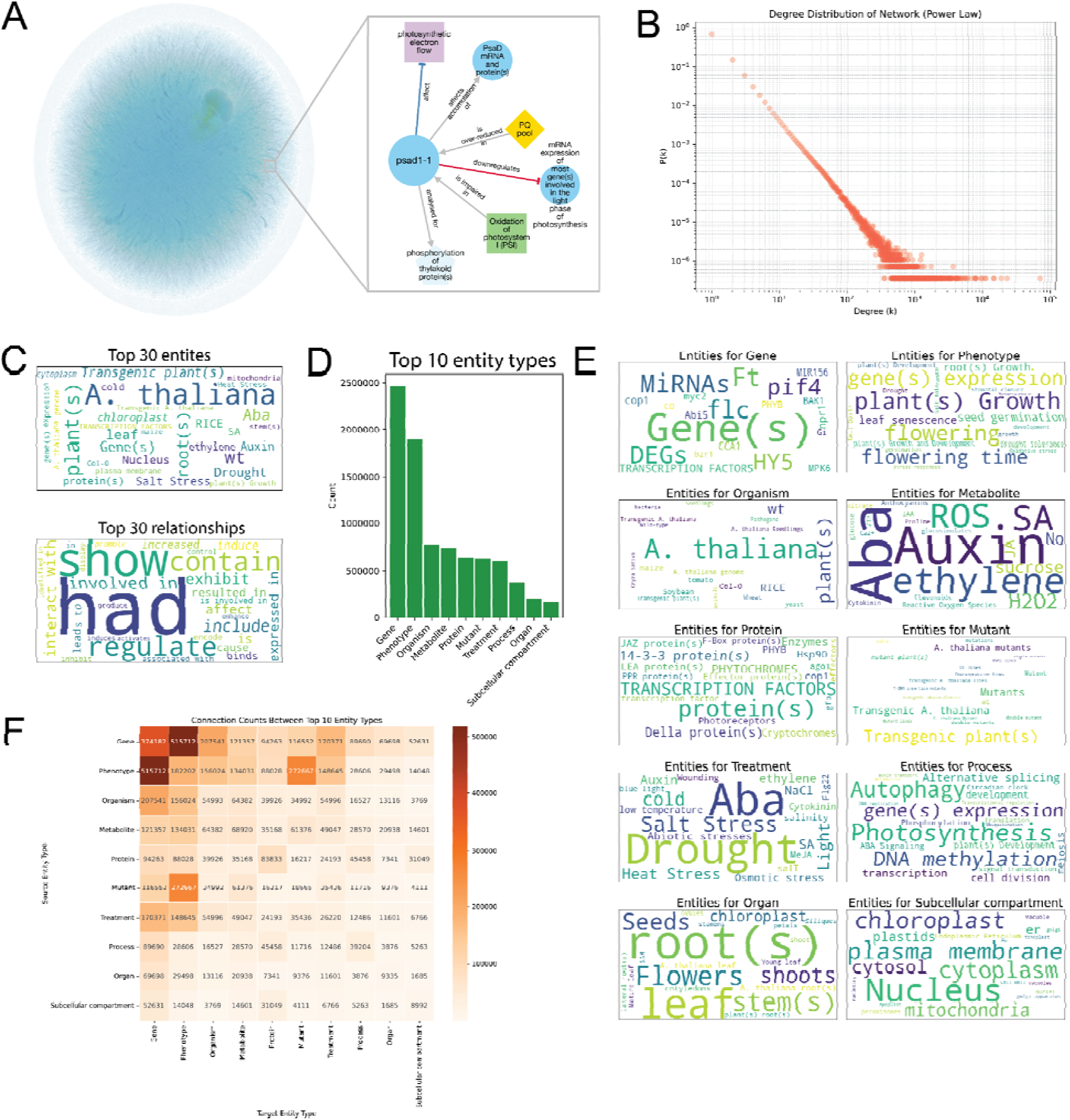
Properties of the Connectome knowledge graph. A) Gephi visualization of the Connectome knowledge graph colored by node degrees. The ForceAtlas 2 algorithm was run until convergence with a stronger gravity law and a scaling factor of 0.5 to separate the graph’s nodes. Light blue nodes represent nodes with the fewest degrees, and light green nodes are the nodes with the highest degrees. B) The degree distribution of the knowledge graph, where nodes represent entities and edges represent relationships between these entities. The x-axis (degree (k)) represents the number of connections (relationships) each entity has. In contrast, the y-axis (frequency P(k)) shows the frequency of an entity having exactly k connections. C) Top most frequently-appearing entities (top) and relationship (bottom) types. The size of the lettering is proportional to the number of relationships. D) Top 10 entity types, with entities (x-axis) and their numbers (y-axis). E) Top 20 entities found in the top 10 entity types. F) The number of edges between the top 10 entity types.

We next investigated which entities and relationships are most important in the KG. General entities, such as ‘A. thaliana’ and ‘plants’, and general relationships, such as ‘had’, ‘show’ were most frequently observed (Figure 3C), but we also observed more specific entities (e.g., auxin, ABA) and relationships (regulate, interacts with). The most common entity types comprised ‘gene’ and ‘phenotype’, in line with our selecting articles containing gene names and ‘*Arabidopsis thaliana*’ (Figure 3D). A closer look at the entity types revealed the most common entities for genes (e.g., *FT, FLC, PIF4*), phenotypes (growth, flowering, germination, gene expression), organism (A. thaliana), metabolite (ABA, auxin, ethylene, ROS), protein (general transcription factors, phytochromes, enzymes), mutants (general ‘transgenic plants’), treatment (drought, ABA, salt stress), process (photosynthesis, DNA methylation, autophagy), organ (root, leaf) and subcellular compartment (nucleus, plasma membrane)(Figure 3E). Finally, we investigated how often the different entity types are connected in the network. We observed most connections between ‘gene’--’phenotype’, ‘mutant’--’phenotype’ and ‘gene’--’treatment’ (Figure 3F), likely reflecting the typical function studies that characterize genes in terms of mutant phenotypes and responses to various treatments.

### Evaluation of the coverage and accuracy of the PlantConnectome

Our main motivation in this study was to expand the amount of the gold standard data capturing experimentally-verified gene functions. We, thus, investigated the overlap of relationships in our KG with data provided in the public repositories.

To compare the coverage and the accuracy of gene regulatory networks (GRNs), we obtained the *Arabidopsis thaliana* gene regulatory network from AGRIS (https://agris-knowledgebase.org/downloads.html, updated March 2019) (Yilmaz et al. 2011), comprising 4,409 confirmed transcription factor -> target edges. We also identified 3,695 edges from a study investigating responses to jasmonic acid (Zander et al. 2020). Next, we identified 15,009 transcription factor -> target edges in the Connectome (Supplementary Data 4). The edges shared between AGRIS and the Connectome comprised of ‘regulate’, ‘binds’, ‘activates’, ‘targets’, demonstrating the Connectome’s ability to identify the various functions of transcription factors (Figure 4A). However, we observed a very minor overlap (e.g., 206 edges between AGRIS and the Connectome) between the three GRNs (Figure 4B), indicating the high dissimilarity between the GRNs. We identified 14,736 Connectome-specific edges between transcription factors and target genes, and while the most frequent association was ‘interacts with’, we also identified typical transcription factor-related terms, such as ‘regulate’, ‘binds’, ‘activates’, ‘represses’, and others (Figure 4C). This indicates that the Connectome transcription factor networks seamlessly integrate protein-protein interactions (PPIs) and GRN networks.

**Figure 4.**
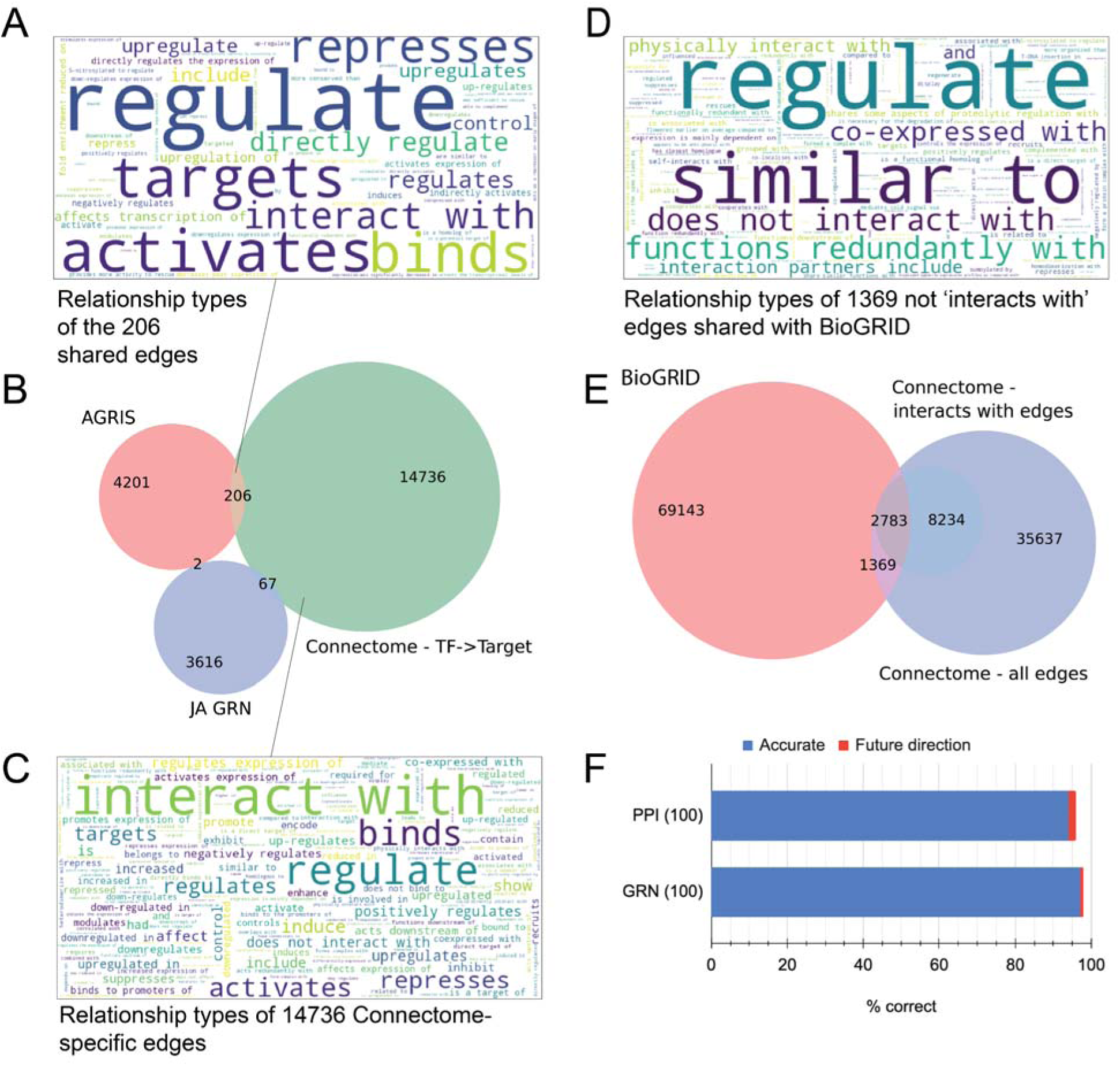
Evaluation of the gene regulatory and protein-protein interaction networks identified by the Connectome. A) Word cloud of the 206 relationship types shared between AGRIS and the Connectome. B) Venn diagram showing the overlap between AGRIS, the Connectome and the Jasmonic Acid Gene Regulatory Network (GRN). The numbers indicate edges found between transcription factors and putative target genes. C) Word cloud of the 14736 Connectome-specific edges. *D) Word cloud of the 1369 edges shared between BIOGRID and the Connectome. The edges do not belong to the ‘interacts with group’. E) The Venn diagram shows the overlap between BIOGRID and the Connectome. All gene-gene edges in Connectome are indicated with a green circle, while the blue circle shows genes connected by ‘interacts with’ relationships.* F) Accuracy evaluation of the protein-protein interaction (PPI) and gene regulatory network (GRN) edges that are specific to the Connectome. The blue and red fields indicate correctly inferred or ‘future direction’ relationships, respectively.

Furthermore, we compared the protein-protein (PPI) network from BioGRID to the Connectome’s network. We found 1,369 edges shared between BioGRID and the Connectome that were not of class ‘interacts with, forms a complex with’ (or similar), and found that the edges comprised of types such as ‘phosphorylates’, ‘associates with’, but also ‘does not interact with’ (Figure 4D). This indicates that the Connectome can provide additional nuances to the PPIs or even contradictory information that should be investigated further. Overall, we also found a relatively poor overlap between BioGRID and Connectome, with only 2,783 ‘interacts with’ edges shared (Figure 4E), and the large majority of edges being specific to each database.

To validate the accuracy of the Connectome-specific edges, we randomly sampled 100 GRN (out of 14,736) and 100 PPI (out of 8,234) edges (code in Supplementary Data 1), and inspected their accuracy by reading the corresponding text (Table S7). Overall, we observed 97% and 94% accuracy for the GRN and PPI networks, respectively (Figure 4F). In agreement with the misclassified entities and relationships (Figure 2B), GPT-4o confused future directions (e.g., ‘In this regard, it will be interesting to investigate whether FUS3 interacts with IDD8 through the fourth ZF domain.) as a fact (‘FUS3 interacts with IDD8’)(Table S7). These results indicate Connectome’s valuable companionship and alternative role to not only AGRIS but also BioGRID.

### Features of PlantConnectome

To provide access to the Connectome, we constructed dedicated database (https://plant.connectome.tools/), which offers numerous methods of searching for genes, metabolites, organs, and other entities by terms, author names, and PubMed IDs, alongside a catalogue page (accessible under the https://plant.connectome.tools/catalogue) listing all entities in the database. An entire information page is also provided for each entity in the connectome, containing its definitions (e.g., CESA: ‘A large family of genes encoding cellulose synthases and related enzymes’) and source article.

To detail PlantConnectome’s search result page, we performed a standard query with the gene “Psad1” (‘Mutant affecting photosystem I complex in plants’, https://plant.connectome.tools/alias/psad1), which is involved in the formation of photosystem I (Ihnatowicz et al. 2004). The entity’s landing page displays the number of nodes in the knowledge graph and the number of papers used to construct the KG, together with the extracted definitions of the entity (Figure 5A).

**Figure 5.**
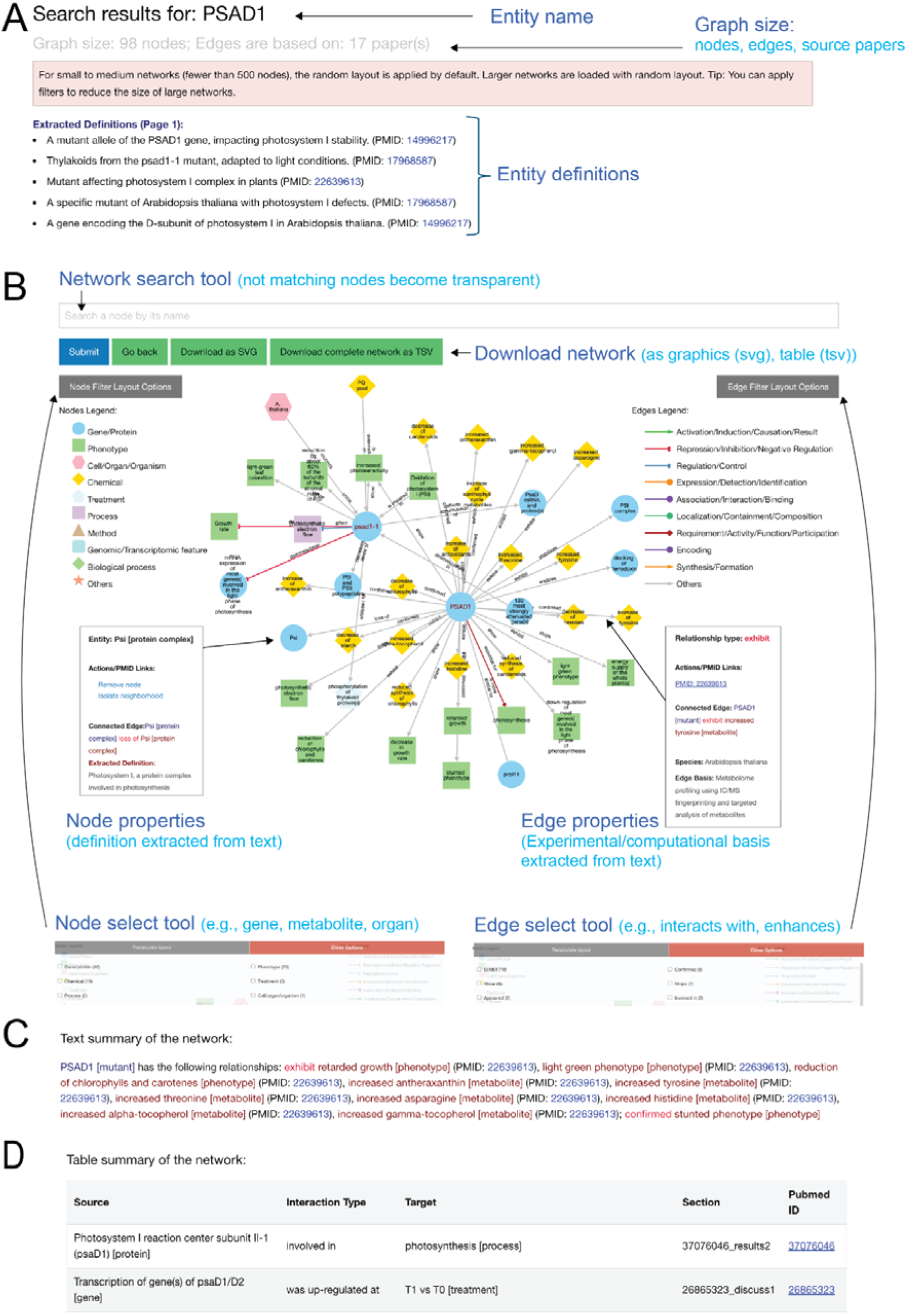
Outline of the Connectome entity page. A) The top of the page contains the entity name and the graph size, comprising the number of nodes (entities) and papers that are used to build the current graph. Note that the graph size changes dynamically when the user adjusts the node and edge selection. The entity definitions extracted from the various papers mentioning the entity are found in the paginated table. B) The graph is visualized as an interactive network implemented in cytoscape.js, where the nodes can be moved around, and the different elements can be clicked on to open tooltips that display additional information. The entities in the network can be searched by writing the entity’s name and clicking submit (blue button). The users can select the entity types (e.g., genes/proteins, metabolites) and edge types (e.g., regulates, interacts with) of interest by clicking on the Node and Edge select tools. The network can be downloaded as a vector graphic image and tab-delimited table. C) The text summary provides an organized, textual representation of the network and PubMed IDs that underpin each edge. The text summary is dynamic and responds to the node and edge selection performed by the user. D) The network is also summarized as a table, where each edge is found as a row in the table.

The knowledge graph is represented as an interactive network, depicting the various relationships the search query shares with other entities in the database (Figure 5B). Upon clicking on a node, the user is provided with a ‘Node properties’ tooltip displaying the node’s definitions, and a set of options enabling the removal of the node or isolation of the node’s neighborhood. Clicking on an edge opens an ‘Edge properties’ tooltip that displays the PubMed ID underpinning an edge and shows the experimental basis (if available) of the edge. Users can select the node and relationship types by clicking on the ‘Node select’ and ‘Edge select’ tools and thus focus on the entity and relationship types of interest (Figure 5B). The current network view can be downloaded as an image (SVG) or as a tab-delimited table, ready for further processing in, e.g., Cytoscape. The network is also available as a text summary (Figure 5C) and a table (Figure 5D), where clicking on a given PubMed ID will prompt a popup containing the corresponding abstract. The text summary adjusts its content to the selections specified in the ‘Node select’ and ‘Edge select’ tools.

Finally, PlantConnectome enables users to perform searches through an API, which returns a JSON object containing relevant network and functional information, extending its functionality to bioinformaticians who desire programmatic access to our database. As an example, an alias search on the PSAD1 gene may be performed by accessing the URL “https://plant.connectome.tools/api/alias/psad1”.

### Examples of how to use PlantConnectome

We provide three case studies, comprising protein complexes, gene regulatory networks, and stress responses, to demonstrate how the Connectome can be used to rapidly summarize available knowledge.

#### Example 1: Chloroplast Protein Translocation and Channel Member TOC75

To exemplify how the Connectome can be used to study protein-protein interactions, we used TOC75 as a query for the ‘alias’ search (https://plant2.connectome.tools/alias/TOC75), which searched for entities labelled as *TOC75* and it’s aliases: *AT3G46740, MAR-01, TOC75-III*. This identified a knowledge graph containing 355 nodes based on 80 papers. We then narrowed down the graph to genes/proteins with the ‘Node selection’ tool, resulting in 199 nodes based on 61 papers.

Translocase complexes on the outer and inner envelope membranes (TOC and TIC, respectively) are used to import proteins into the chloroplast (Stengel et al. 2009). We compared the TOP75 graph to a review on the translocase complexes (Richardson and Schnell 2020), which revealed known TOC75 interactions such as *TOC 22, 34*, *159*, and *TIC236* (Figure 5A). The associated nodes also provide additional genes relevant for TOC75 function, such as dek5 mutant, which is reducing the levels of TOC75 (experimental organism: maize, evidence: proteomics analysis and immunoblotting of chloroplast envelope proteins)(Zhang et al. 2019), and chaperone HSP90C that interacts with many of the translocon proteins (experimental evidence: Coprecipitation experiments with protein import components)(Inoue et al. 2013). Thus, the Connectome allows a rapid elucidation of protein-protein interactions, and provides source literature and evidence types supporting these interactions.

#### Example 2: Secondary Cell Wall Master Regulator

To demonstrate how our database can be used to study gene regulatory networks, we selected the secondary cell wall biosynthesis regulator, *MYB46* (https://plant.connectome.tools/alias/MYB46). The initial network contained 618 nodes based on 129 papers, but we selected edges capturing typical gene regulatory relationships (e.g., ‘regulate’, ‘directly activate’) from the ‘Edge filter’ menu, and arrived at a network comprising 211 nodes based on 75 papers (Figure 6B). A literature search on the gene regulatory network underlying secondary cell wall formation revealed a large overlap between the output of the Connectome and the figure in the review article (Xiao et al. 2021).

**Figure 6.**
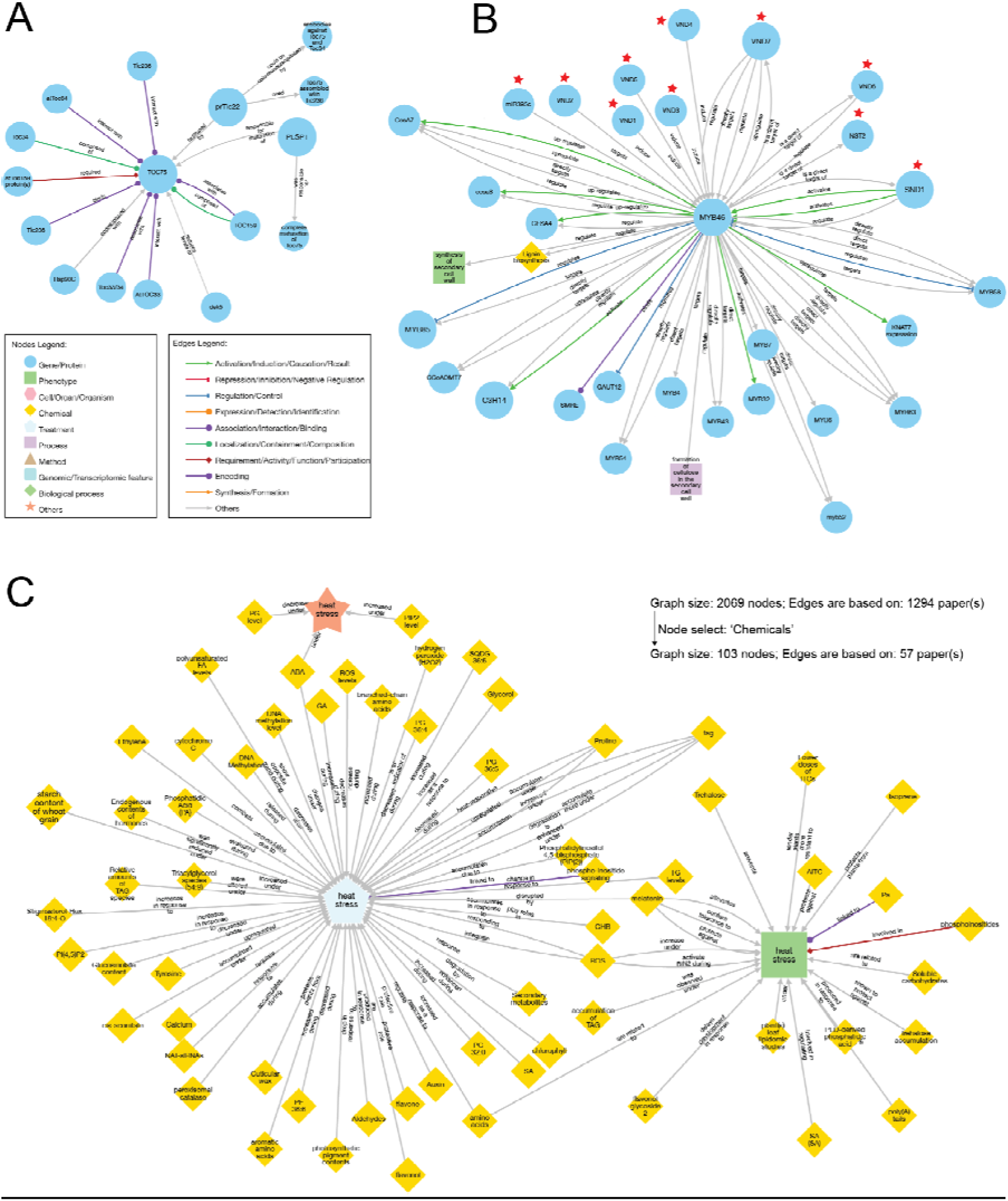
Usage examples of the Connectome database. A) Gene/protein neighborhood of TOC75 Translocon at the outer envelope-membrane of chloroplasts 75 protein, found by alias search using ‘TOC75’. Nodes represent gene/protein entities, while edges depict relationships. The node and edge legend indicate the different types of entities and relationship types, respectively. B) Gene/protein neighborhood of MYB46. Transcription factors regulating MYB46 are indicated with red stars. C) Neighborhood of ‘heat stress’ entities. The nodes were selected to only comprise of ‘chemical’ entity type.

Genes regulated by *MYB46* included the secondary cell wall cellulose synthases (CESA4,7,8), lignin biosynthesis genes, and a panel of downstream transcription factors (*MYB4, 7, 32, 43, 52, 54, 58, 63*). Genes regulating *MYB46* comprised transcription factor *SND1* and micro-RNA miR395c (Figure 6B). Interestingly, our Connectome, but not the review, identified *VND1-7* and *NST2* (evidence: Direct target analysis using the estrogen-inducible system)(Zhong et al. 2008) and *VNDs* (evidence: Quantitative PCR analysis of *VND1* overexpressors showing induction of *MYB46* gene expression)(Zhou et al. 2014) as regulators of *MYB46*. On the other hand, our Connectome missed downstream *MYB20, 42, 75, 103* (Xiao et al. 2021).

Since the KnowledgeNetwork is summarized in a human- and machine-readable format, we investigated whether ChatGPT can generate publication-ready passages. To this end, the network summary (exemplified in Figure 5C) was fed into ChatGPT-4o with the following prompt:

*‘Summarize the function of MYB46, using the text below. Indicate which genes it regulates. Indicate which genes regulate it. Indicate which processes it regulates. For each statement, make sure to indicate the PubMed ID, for example: ‘Gene A regulates gene B (PubMed ID)’. Use only the information provided by this text:*

*<NETWORK summary text pasted in here>*

*Write the text as if it was a paragraph for a review paper. List ALL of the genes mentioned in text.’*

The outcome is, in our opinion, on par with some review articles:

##### 1) Function of MYB46

MYB46 (At5g12870) is a transcription factor that functions as a master regulator in secondary cell wall (SCW) biosynthesis (PMID: 24984711, 24009617). It directly regulates the expression of genes involved in the biosynthesis of cellulose, hemicellulose, and lignin (PMID: 24984711, 34493336). MYB46 plays a key role in activating the secondary wall biosynthetic program (PMID: 25148240), promotes SCW thickening (PMID: 33086706), and is crucial for secondary wall formation in fibers and xylem vessels (PMID: 25535195, 34879821). Furthermore, MYB46 has been shown to drive differentiation of xylem vessel cells (PMID: 34971462), enhance resistance to Botrytis cinerea (PMID: 32214345), and regulate plant immune response (PMID: 34208611). The protein also acts in abiotic and biotic stress responses, including its upregulation under stress (PMID: 31077628, 33826618). MYB46 functions in a transcriptional regulatory cascade (PMID: 19808805) and activates transcription of lignin, cellulose, and hemicellulose biosynthetic genes (PMID: 33537043). Its overexpression can induce ectopic secondary wall biosynthesis (PMID: 34493336), leading to stunted growth (PMID: 25535195).

##### 2) Genes Regulating MYB46

MYB46 is regulated by several transcription factors. It is a direct target of SND1 (secondary wall-associated NAC domain protein 1), NST1, NST2, VND6, and VND7, which are part of the NAC transcription factor family involved in secondary wall biosynthesis (PMID: 37372306, 29973639). The VND family members (VND1-7) also induce the expression of MYB46 (PMID: 25148240, 35154217). Additionally, third-tier SWNs (NAC TFs) regulate MYB46 (PMID: 24009617), while MPK6-mediated phosphorylation is involved in post-translational regulation of MYB46 (PMID: 33826618). MYB46 is upregulated under stress by various stimuli (PMID: 33826618), and SnRK2.2/3/6 kinases regulate its expression during stress responses (PMID: 33495344). GbTCP5 (PMID: 34880892) and ERF041-VP16 (PMID: 33995450) are also involved in the transcriptional regulation of MYB46.

##### 3) Genes Regulated by MYB46

MYB46 regulates a wide array of downstream genes involved in secondary wall biosynthesis. These include cellulose synthase genes such as CESA4, CesA7, and CesA8 (PMID: 23726771, 26011122, 37372306), as well as xylan and lignin biosynthetic genes (PMID: 34493336, 24885077). It activates the expression of other transcription factors like MYB58, MYB63, MYB85, and MYB43, which further control SCW biosynthesis (PMID: 24009617, 31881731). MYB46 also regulates genes involved in hemicellulose synthesis, such as AtCSLA9 and GAUT12 (PMID: 24243147, 25120548). Additionally, it directly activates secondary wall biosynthetic genes (PMID: 22197883, 34862967), including mannan synthase and other SCW-associated enzymes (PMID: 24879533, 32775951). MYB46 also influences genes linked to lignin biosynthesis, such as CSE and CCoAOMT7 (PMID: 27390589, 35154217), and promotes SCW-related gene expression (PMID: 36077531).

Importantly, since ChatGPT was asked to use the existing PubMed IDs, it does not hallucinate non-existing references.

#### Example 3: Heat stress

To demonstrate how the Connectome can be used to study entities that are not necessarily genes, we investigated chemicals, hormones and metabolites involved in heat stress. We searched for ‘heat’ with ‘normal’ search, which took us to the page of entities containing ‘heat’ in their description (https://plant2.connectome.tools/normal/heat). The resulting large knowledge graph comprised 2069 nodes from 1294 papers, which is expected as heat stress is one of the most studied abiotic stresses in plants (Koh et al. 2024). We further focused the search by selecting ‘Chemical’ in the ‘Node filter’ tool, which shrank the graph to 103 nodes based on 57 papers (Figure 6C). This revealed two central ‘heat stress’ nodes, categorized as ‘phenotype’ and ‘treatment’. The graph revealed multiple chemicals that are important for heat stress tolerance, such as isoprene (found in Discussion in (Weraduwage et al. 2023), evidence: Transcriptomic studies on Arabidopsis thaliana fumigated with isoprene), trehalose (introduction in (Jin et al. 2016), no evidence available), AITC (Allyl isothiocyanate, found in Introduction of (Øverby et al. 2015), no evidence available) and flavonols and flavones (found in Discussion in (Liu et al. 2021), evidence: LC-MS measurements on QE HF-X coupled to Vanquish UHPLC). The graph also reveals compounds that change their levels under heat stress, such as triacylglycerols (Introduction of (Higashi et al. 2015), evidence: LC-MS-based lipidomic analysis), phosphatidic acid (Discussion in (Kocourková et al. 2021), no evidence available), reactive oxygen species (ROS, Discussion in (Cocetta et al. 2022), evidence: histochemical analysis using 3,3-diaminobenzidine (DAB) staining and literature references). The graph also mentions several hormones, such as salicylic acid (SA), auxin and ethylene, cellular structures such as cutin and starch content, and other entities that were classified as chemicals by GPT-4o. To conclude, the dynamic selection option of edge types in the network enables scrutinizing different relationship types between the entities found in PlantConnectome.

## Discussion

We have illustrated GPT’s text mining capacities in the context of scientific literature, processing over 71,000 research abstracts at a moderate cost (∼5000 USD) and harvesting invaluable functional information therein. GPT could extract key entities and relationships from research paper abstracts with high accuracy (Figure 2B, 4E) and few prompts (Table S5). The amount of functional information excavated from the abstracts increased the amount of machine-readable data, as demonstrated by our gene regulatory networks that nearly tripled the available data (Figure 4B). Moreover, PlantConnectome overcomes the limitations of typical databases that employ only one data type, as it draws upon numerous data sources in establishing gene functions, organ development, gene regulatory networks, protein-protein interactions, and other phenomena, all in a user-friendly manner.

Our evaluation has shown that PlantConnectome is not only comprehensive and accurate but also complementary to existing databases (Figure 4). Comparing PlantConnectome’s gene regulatory networks against AGRIS and its protein-protein interaction networks against BioGRID demonstrates that PlantConnectome’s retrieved networks do not largely overlap with these reference databases. Rather, the GPT-extracted networks complement them, showing the effectiveness of our text-mining approach in utilizing the vast amount of literature that has not been captured by manual curation.

However, GPT’s outputs are not entirely accurate and still warrant manual verification, as GPT-4o models have a tendency to misidentify entities and relationships (Figure 2B, 4E), which is perhaps attributable to the varying language and content of the >71,000 processed articles. *The correction of errors may be carried out by fine-tuning the models with manually curated examples containing the expected output (as, for instance, that found in Table S6). Thus, the users of our database and knowledge graph are encouraged to click on nodes and edges to further validate these entities’ accuracy*.

In conclusion, PlantConnectome is an innovative tool, combining the power of a state-of-the-art language model with the comprehensive information embedded in a massive collection of research articles. The tool offers an efficient and diversified way to retrieve information for genes, metabolites, tissues, organs, and other biological components. The potential applications of PlantConnectome are wide-ranging and extend beyond those we have highlighted in this article. Furthermore, since we only analyzed articles mentioning *Arabidopsis thaliana* and it’s genes, the inclusion of all plant scientific literature together with the inclusion of more full-text papers is bound to increase the completeness of the knowledge graph, help us stay up to date with the plant literature, and provide gold standard data for gene function prediction studies. We anticipate that PlantConnectome will become a valuable resource for the plant science community to facilitate various research activities, from a preliminary investigation of gene functions to an in-depth study of a particular biological process.

## Supplemental Figures

**Figure S1.**
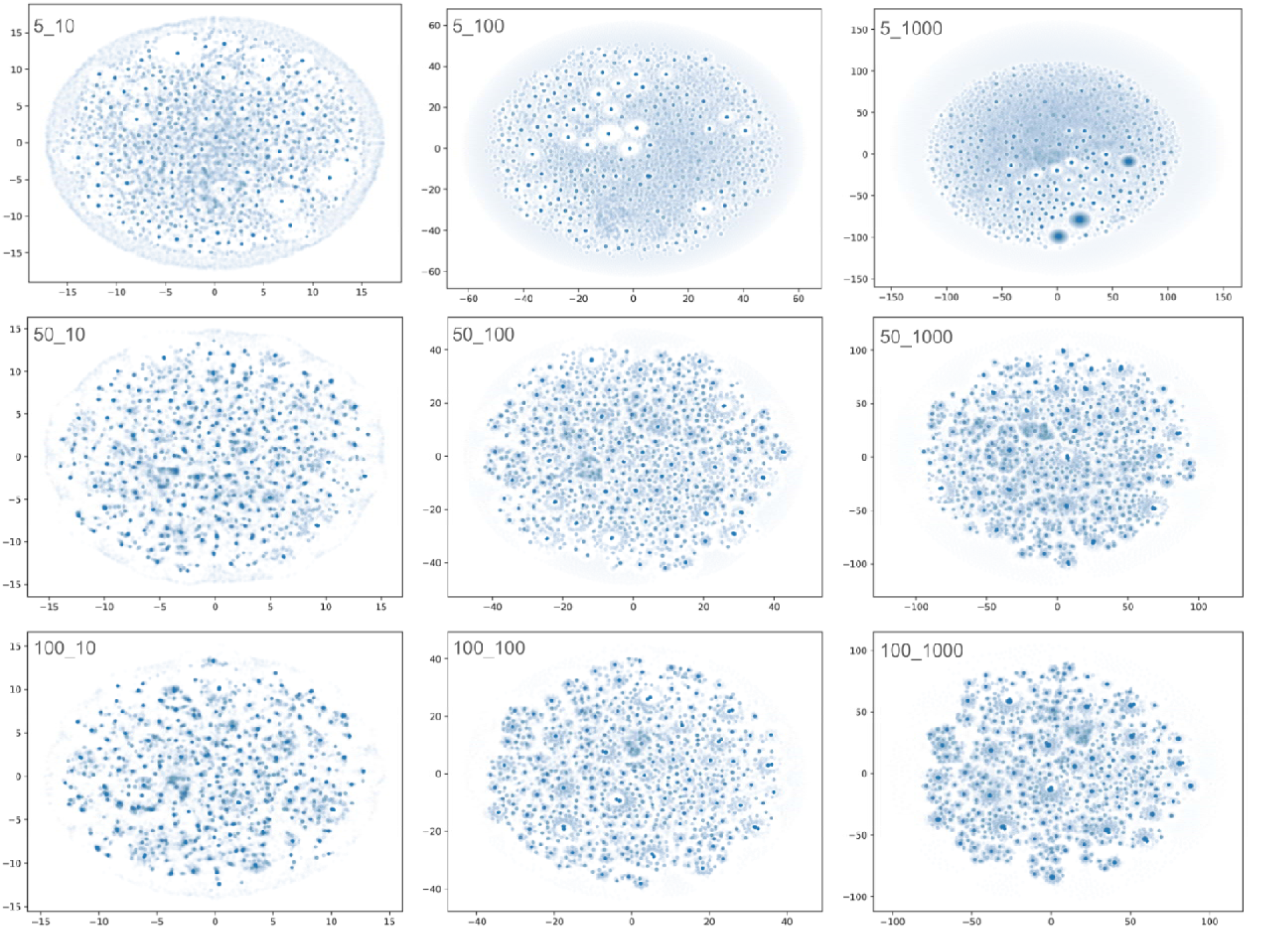
tSNE analysis of the abstracts at the different perplexity and iteration values. The evolution of the plot at a perplexity of 5 (first row), 50 (second row) and 100 (third row) and different ranges of iterations: 10 (first column), 100 (second column) and 1000 (third column).

**Figure S2.**
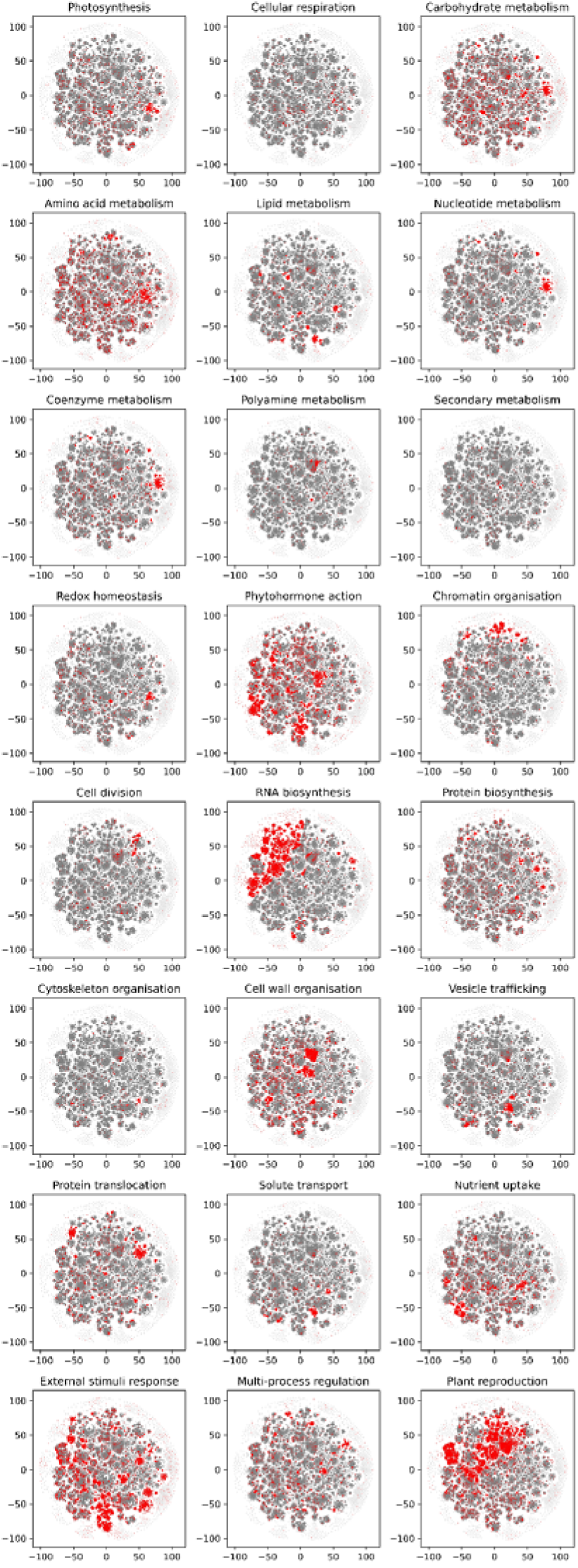
tSNE analysis of the abstracts of the different biological processes, as defined by MapMan. A red point indicates an abstract that contains a keyword (e.g., pollen is a keyword for plant reproduction), while grey point indicates an absence of the keyword match.

**Figure S3.**
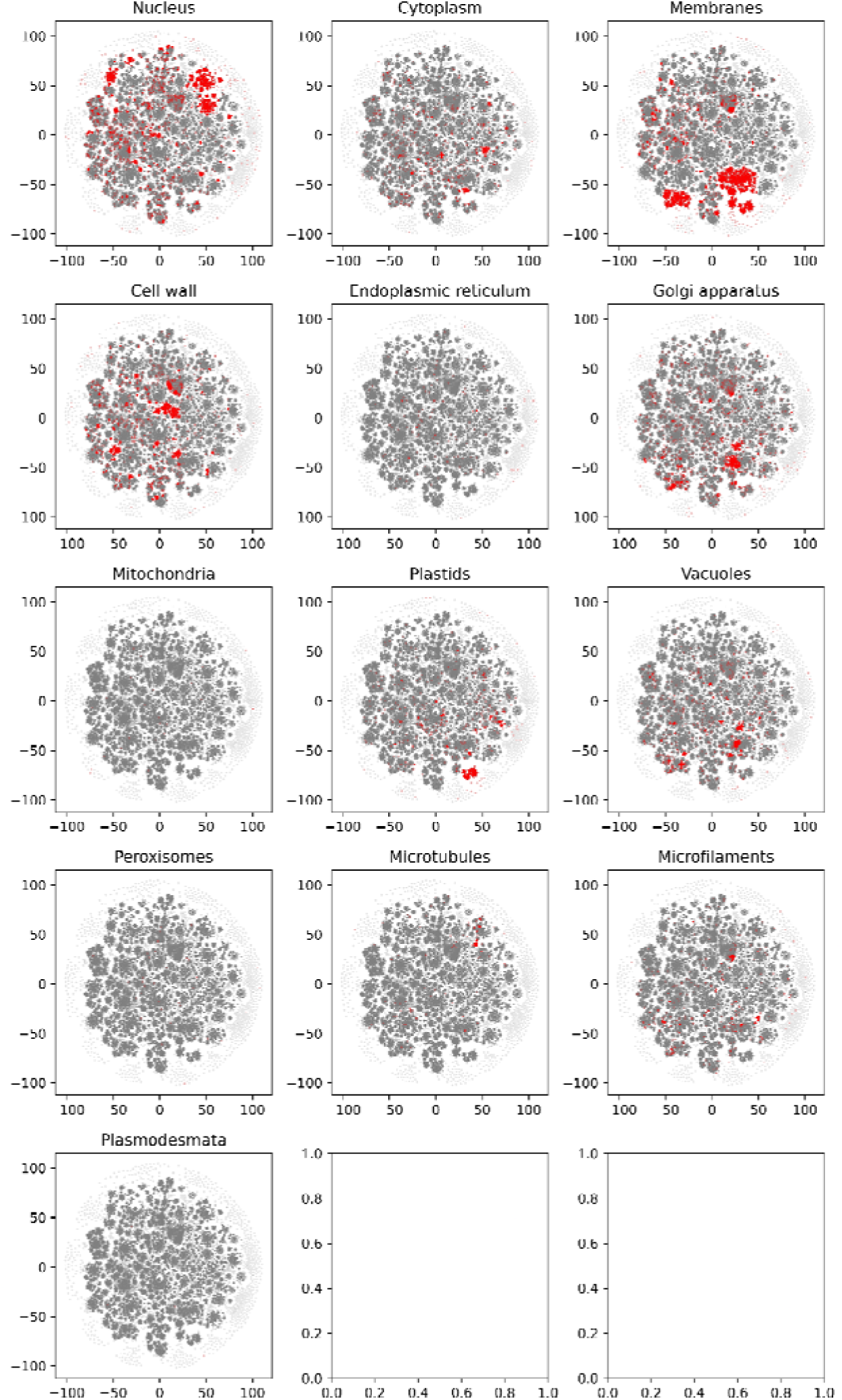
tSNE analysis of the abstracts of the different cellular compartments.

**Figure S4.**
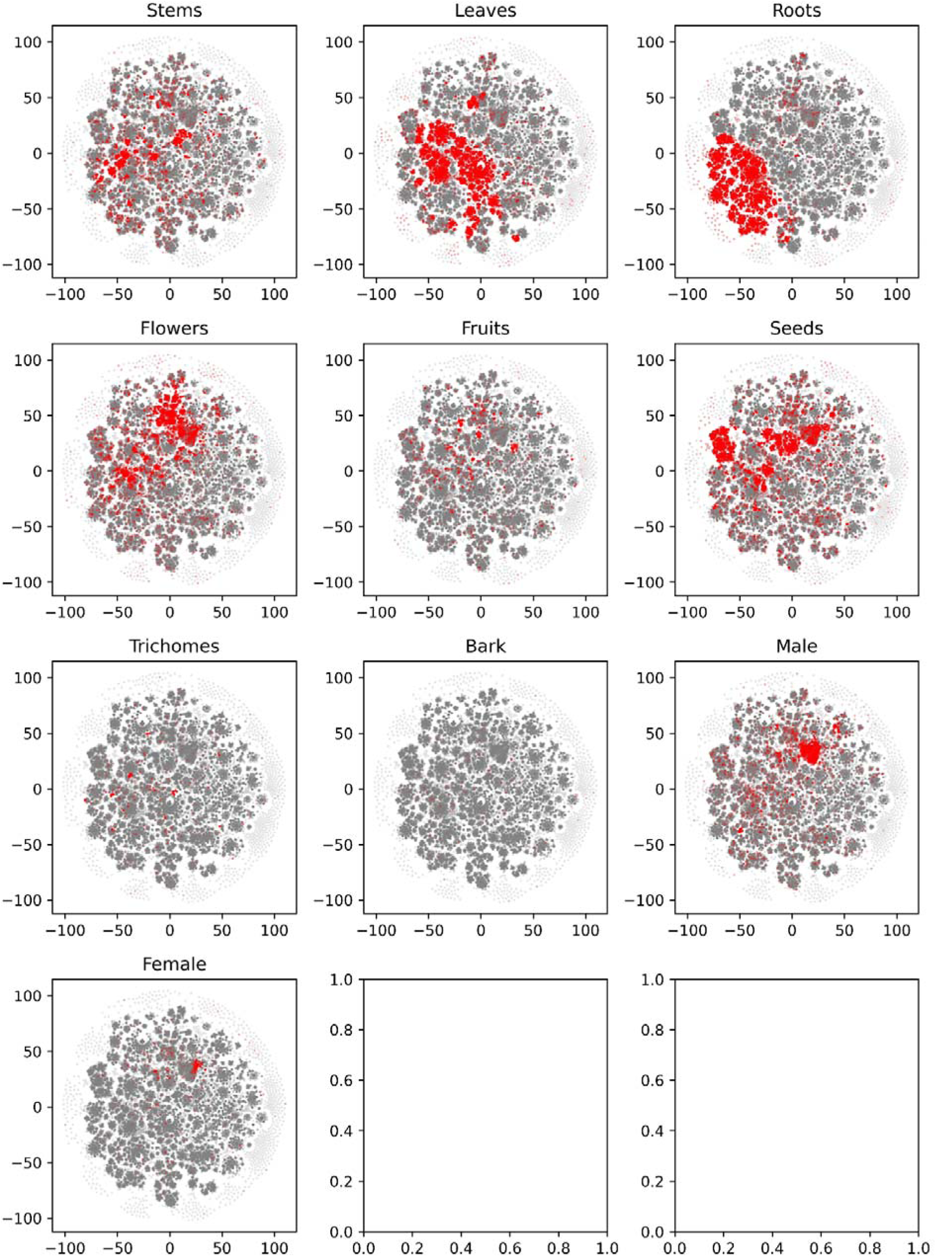
tSNE analysis of the abstracts of the different major organs and cell types.

**Figure S5.**
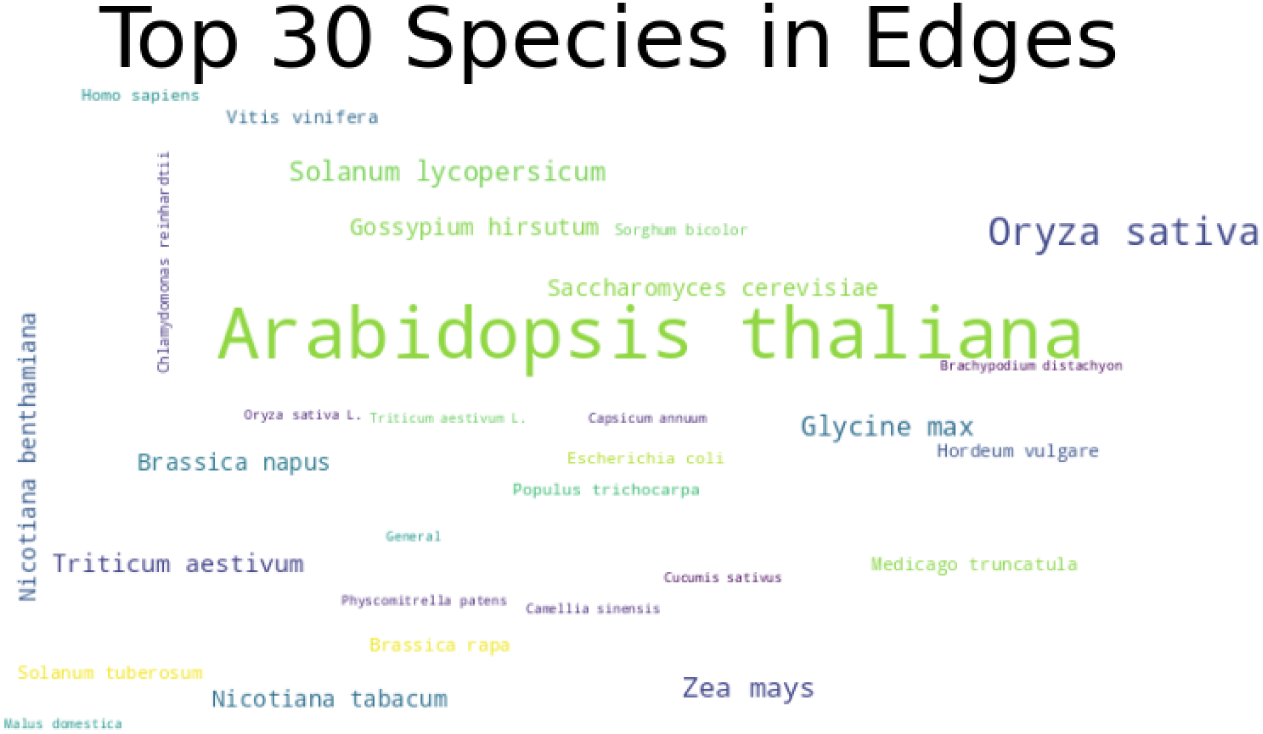
Wordcloud depicts the species in which the relationships (edges) were reported in.

## Supplemental Tables

**Table S1. Gene IDs (column A) and aliases (column B) used to query the NCBI database for articles.**

**Table S2. Journals used to construct the database.** The columns indicate the journal name, article title, publication year, PubMed ID, and whether the article was analyzed as full text (yes/no).

**Table S3. The three categories, their sub-categories, and keywords used to perform topic analysis of paper abstracts.**

**Table S4. Keyword co-occurrence analysis.** The two keywords are in columns A and B, and the number of articles in which these keywords co-ocurred are shown in column C.

**Table S5. The prompts used to build the knowledge graph, extract entity definitions and edge basis.**

**Table S6. Edge and entity type accuracy evaluation.** Each row represents and edge in the knowledge graph. Each edge contains the PubMed ID, source and target nodes, source and target entity types and relationship description. The evaluations of the source types (column G), target types (Column J), and relationship (Column M) are indicated. The comments and the sentences that underpin the edge are found in the other columns.

**Table S7. Evaluation of the accuracies of the gene regulatory edges (top sub-table) and protein-protein interactions (bottom sub-table).**

## Supplementary Data

**Supplementary Data 1. The code to download and process the papers, and generate the figures in the manuscript is available at** https://colab.research.google.com/drive/1gENxLK2172Bq1sV_dO6JhvORsFgaq0fS?usp=sharing

**Supplementary Data 2. Keywords present in each abstract analyzed in this study.** The keywords are defined in Table S3. https://figshare.com/ndownloader/files/49392538

**Supplementary Data 3. Co-occurence network of keywords found in each abstract.** The network can be viewed in Cytoscape. https://figshare.com/ndownloader/files/49392595

**Supplementary Data 4. The knowledge graph used to build the Plant Connectome.** The edges and nodes have been disambiguated by the code found in Supplementary Data 1. https://figshare.com/ndownloader/files/49198933

## Data availability

The Plant Connectome database source code is available at: https://github.com/mutwil/plant_connectome_latest/

